# Comparison of antibody titres between intradermal and intramuscular rabies vaccination using inactivated vaccine in cattle in Bhutan

**DOI:** 10.1101/499145

**Authors:** Karma Wangmo, Richard Laven, Florence Cliquet, Marine Wasniewski, Aaron Yang

## Abstract

In developing countries, the cost of vaccination limits the use of prophylactic rabies vaccination, especially in cattle. Intradermal vaccination delivers antigen directly to an area with higher number of antigen-presenting cells. Therefore, it could produce equivalent or higher antibody titres than conventional intramuscular vaccination even when a lower dose is given.

This study aimed to compare the antibody response in cattle vaccinated intramuscularly with 1mL of inactivated rabies vaccine (Raksharab, Indian Immunologicals) against intradermally vaccinated cattle with 0.2mL of the same vaccine. The study was conducted in Haa province of Bhutan where rabies is not endemic. One hundred cattle from 27 farms were selected for the study. Virus neutralising antibody (VNA) response was measured using the fluorescent antibody virus neutralisation test on the day of vaccination (day 0) and 14, 30, 60 and 90 days later.

Overall, 71% of intradermally vaccinated cattle and 89% of the intramuscularly vaccinated cattle produced a protective response (≥0.5IU/mL). This difference was significant (P<0.02) on days 14 and 30 post vaccination with 36 and 58% in the intradermal group having titres ≥0.5 IU/mL respectively compared to the equivalent figures of 78 and 77% in the intramuscular group. The mean VNA titres were lower for intradermal group than intramuscular group (p<0.001) with the mean difference being greater than 0.6 IU/mL. Although low dose intradermal vaccination did produce a detectable antibody response, it was inferior to intramuscular vaccination. Thus, although intradermal vaccination has the potential to reduce the cost of vaccination by reducing the dose required, this study showed that a single dose of 0.2mL intradermally was inferior to an intramuscular dose of 1mL. Further research evaluating dose and dose regimen is needed before intradermal vaccination using the Raksharab rabies vaccine can be recommended in cattle.

## Introduction

Rabies is a fatal zoonotic viral disease inducing an acute disease of the central nervous system in almost all mammals (1). In cattle, rabies causes significant economic losses to livestock farmers (2-5) principally through increasing mortality, decreasing bodyweight and milk production. Furthermore, costs of euthanasia, diagnosis, replacement and vaccination of at risk herds add to the loss (6, 7).

In Bhutan, rabies is endemic in 29% of the country (8) with the dog being the principle vector and reservoir host. Cattle contract the disease as a spill-over infection from dogs; nevertheless over 80% of the economic loss due to rabies in Bhutan is due to cattle deaths (9). The extensive grazing system (>60% of farms (10), free movement of cattle around their compound (>70%) and limited restrictions on access even when they are housed increase the risk of cattle contracting rabies from rabid dogs (11). This risk is exacerbated by the large number of dogs in Bhutan (dog population estimated at 120 000 dogs, equivalent to one dog for every 2.5 cattle) of which 40% are stray dogs, and 31% owned but free roaming (12). Mass vaccination against rabies and sterilization of dogs have been carried out throughout the country on an annual basis to reduce the risk of rabies. However, preventive control measures in cattle, especially vaccination, have been a lower priority, because the risks of virus transmission through cattle are low (13). Nevertheless, in endemic areas of Bhutan, rabies remains common in cattle and causes considerable economic losses to smallholder cattle farmers. Until rabies is eliminated in the reservoir hosts (principally dogs), free movement of reservoir hosts across the India/Bhutan border and from rabies endemic to non-endemic areas combined with limited and accessible housing of cattle means that rabies will continue to be a major concern in Bhutanese cattle with a significant economic issue for the individual affected smallholders.

As economic constraints at both national and local levels are key drivers for the lack of prophylactic vaccination in Bhutanese cattle, finding a way to reduce the costs of bovine vaccination could be useful for initiating stakeholders to improve vaccination coverage of cattle against rabies without markedly increasing the overall cost of rabies control. One potential method of reducing the cost of vaccination is to use intradermal vaccination. Compared to subcutaneous or intramuscular vaccination, intradermal vaccination results in the direct stimulation of a large population of active antigen-presenting cells (14). This could, potentially, increase the magnitude of the immune response even if vaccination is used at a lower dose than that required for the intramuscular route.

Lower dose intradermal vaccination against rabies has been shown to be effective in humans, laboratory animals and dogs (15-17). Data on use of intradermal rabies vaccination in cattle are sparse. A study by Koprowski et al. (18) reported that cattle (11 animals) vaccinated intradermally in the neck region with an attenuated rabies vaccine (1mL) produced protective antibody titres 30 days after vaccination. Asokkumar et al. (19) reported that intradermal vaccination of cattle with an inactivated vaccine (at 1/10 dose) was as effective at stimulating virus neutralising antibody (VNA) titres as the standard intramuscular dose. However, this was a post-exposure study and each animal received multiple doses of vaccine, so it has limited utility in determining the value of preventive intradermal vaccination. Benisek et al. (20) compared intradermal and intramuscular vaccination in a pre-exposure prophylaxis in cattle and found that intradermal vaccination (at 1/5 dose) produced higher mean VNA titres than the intramuscular route. This study had a small sample size (only 10 animals per group) and a limited statistical analysis. In addition, the variance in VNA titres of cattle treated using intradermal vaccination was much higher than that of cattle vaccinated intramuscularly. Thus, although these studies support the hypothesis that intradermal rabies vaccination could be effective in cattle, they do not provide sufficient proof to recommend it for preventive vaccination of cattle during mass parenteral vaccination programmes.

The aim of this study was to compare rabies VNA titres produced by intradermal vaccination at 1/5 of the recommended intramuscular dose (i.e. 0.2 mL) against 1 mL of intramuscular vaccination using the same vaccine in cattle in Bhutan.

## Materials and Methods

All animal use was approved by the ethical committee of the research and extension division of the Bhutan Department of Livestock dated 10^th^ March 2017 and in accordance with the requirements of the Research Code of Practice for the Care and Use of Animals for Scientific Purposes.

### Study design

The study was a multi-site non-inferiority trial with animals randomly allocated to either intradermal rabies vaccination using 1/5 of the recommended dose or standard intramuscular rabies vaccination.

### Study area

The study was conducted in Haa district, which is located in north-western part of Bhutan. As of 2016, the district had approximately 9119 cattle and 1031 farms or household with cattle. Herd size ranged from one to hundred cattle (10). As of March 2017, there had been no record of rabies in either dogs or cattle in Haa for five consecutive years (personal communication, Veterinary Officer, Haa). However, there are risks of future outbreaks as this district shares borders with other rabies endemic districts.

### Sample size calculation

Sample size was calculated using the power calculation for a continuous outcome non-inferiority trial (https://www.sealedenvelope.com/power/continuous-noninferior/). At the 95% significance level with 0.63 as the standard deviation of the outcome (from (20)) and a non-inferiority limit of 0.5, 45 animals were required in each group to detect if there was truly no difference between the intramuscular and intradermal route of vaccination in eliciting protective rabies virus neutralising antibody titres in the vaccinated cattle. The non-inferiority limit of 0.5 was chosen considering the threshold limit for protective rabies antibody titres of 0.5 IU/mL, and an expectation that peak titres produced by intramuscular vaccination would be at least 1.0 IU/mL.

### Farm selection

The district annual livestock statistics records were used to select the animals for this study. All the data were recorded in an Excel sheet (Microsoft, USA). Of the six sub districts, three were excluded as they practiced the transhumant system of rearing, which meant that animals would not be available for follow up. The remaining three sub-districts, which as of 2016, had 523 farms and 3312 cattle were included in the study. All farms recorded as having less than four cattle in the herd were excluded by manual selection, leaving 271 farms. Twenty-five farms were selected randomly from this list using a random allocation table. Based on the census, these 25 farms had 260 cattle with maximum herd size of 50.

### Animal selection

A minimum of four cattle per farm was required to get a final sample of 100 animals from 25 farms. However, on the day of vaccination, 12 of the pre-selected farms had less than four cattle in their herd, so extra farms that were located near to the pre-selected farms were used in addition to the original 25 farms to reach the target of at least 90 vaccinated cattle. In addition, on four farms more than four cattle were selected as four of the pre-selected farms practised the transhumant system of rearing. Only animals of age six months and above were eligible for selection in order to avoid any interference from maternal antibodies. On each farm, cattle were randomly assigned to treatment using a random allocation table. Ninety animals were designated for treatment ‘A’ or treatment ‘B with 45 cattle in each treatment group. Ten cattle were kept as controls.

The age, body condition score (BCS) (0-5 scale) (21) and breed of the selected cows were recorded. Age ranged from 6 months to 15 years, BCS from 2-3, and the selected animals included a mixture of local, Jersey, Jersey cross and Holstein Friesian breeds.

### Vaccination

Selected cattle were given either 1 mL of rabies vaccine (22) intramuscularly into the middle third of the neck, or 0.2 mL of the same vaccine intradermally at the same site. The vaccine used was an inactivated cell culture vaccine authorised for intramuscular or subcutaneous use in cattle, dogs and cats (Raksharab company data). Potency testing at the OIE/EU/WHO reference laboratory on Rabies, Nancy, France showed that the vaccine used in the study had a potency of 8.5 IU/mL (Limits 2.2 to 37.4 IU/mL), greater than the recommended minimum of 1 IU/dose (23, 24).

On farms that had more than four cattle a control cow was selected for blood sampling but not vaccination. These control animals were equally distributed across the three selected sub districts.

All animals, irrespective of treatment group were managed as normal by the owners who were blinded to treatment allocation. Cattle on 17 of the 27 farms were housed permanently during the winter and stall fed; on 10 farms cattle were sent for grazing during the daytime in the winter. On 26/27 farms, during the summer, all adult cattle were released for grazing during the daytime and housed only at night. One farm housed cattle throughout the year, using two different sheds, one for winter and one for summer. The study started in late winter (March) and continued through to summer (June). Day 0 sample collection and vaccination was undertaken on 9^th^ to 11^th^ of March. On most farms, animals were housed for the first three sample collections (days 0, 14 and 30), but on days 60 and 90 all farms had started summer grazing except for the farm which housed their cows throughout the year.

### Sample collection

Blood samples were collected on day 0 before vaccination and on days 14, 30, 60 and 90 after vaccination. To avoid grazing times, samples were collected from 6 am to 11 am and from 4 pm to 8 pm; approximately 10 mL of blood was collected via jugular venepuncture into plain vacutainers (BD, India). After collection, blood samples were allowed to settle for 10 to 15 minutes before moving on to the next farm.

For the blood samples collected in the morning, serum separation was undertaken during the afternoon (1 pm-3 pm), while for those collected in the evening, serum separation was done at night (9 pm-11 pm). Blood samples were centrifuged at 1000g for 15 to 20 minutes, and serum transferred to screw-capped cryovials for storage at −20°C (25). Duplicate aliquots were collected for each sample. Each sample from a cow was assigned a sample identification number (order of sampling) so that the laboratory was blinded to the treatment group.

At the end of the sample collection period (i.e. late June), one serum aliquot per cow was transported by air to the OIE/EU/WHO reference laboratory on Rabies, Nancy in France for analysis.

### Seroneutralization test

The serum samples were analysed for VNA titres using the standard fluorescent antibody virus neutralisation test (FAVN) protocol as described by Cliquet et al. (26). Briefly, the test was performed on 96 wells microtiter plate with serial threefold dilutions of the positive, negative controls and the test sera. The Challenge Virus Standard-11 ATCC number: VR 959, rabies virus produced in BHK-21 cell lines (ATCC number: CCL-10) was used as the challenge virus and Dulbecco’s modified Eagle’s medium with 10% fetal calf serum and antibiotics as a diluent.

50 μL of diluted challenge virus containing around 100 TCID_50_ was added into the wells containing diluted samples and diluted positive and negative controls. A back titration of the diluted challenge virus was also performed. The microplates were then incubated at 36°C+/- 2°C in a 5% carbon dioxide (CO2) humidified incubator for 1 hour. Following incubation, 50 μL of diluted BHK-21 cell suspension with concentration of 4 × 10^5^ cells/mL was added to each well and further incubated at 36°C±2°C in a 5% CO2 humidified incubator for 48 hours. The cell culture medium was then discarded and the microplates rinsed once in phosphate buffer solution and then in 80% acetone. The microplates were then fixed in 80% acetone at room temperature for 30 minutes which was drained off and air dried at room temperature.

Fluorescein isothiocyanate conjugate was used for staining microplate wells. The wells were read using a fluorescent microscope (between 100X and 125X magnifications) and examined for the presence or absence of fluorescent cells. It is an “all or nothing” reading therefore the well was considered positive if at least one fluorescent cell was detected.

### Data analysis

Three thresholds of antibody titre were used to indicate response to vaccination: ≥0.17 IU/mL (minimum titre; (24)), ≥0.24 (potentially protective titre; (27-29) and ≥0.5 IU/mL (indicative protective titre; (23, 30). The proportion of cattle meeting these thresholds at each time point were compared for the two treatment groups using a generalised linear model with a binomial output and a logit link, with cow as a random effect and time since vaccination and treatment group (and their interaction) as fixed effects.

The effects of time and treatment group on antibody titre were then analysed. The titre data were significantly right skewed so were log transformed before analysis. A repeat measures mixed model was used with log VNA titres as the outcome variable, cow as a random effect and time since vaccination and treatment group (and their interaction) as fixed effects. A heterogeneous first-order autoregressive covariance structure was used based on minimising the Akaike information criterion. These analyses were undertaken using SPSS statistics 24 (IBM, USA).

Further analysis was then undertaken to include factors other than treatment and time, and to establish; 1) variables that affected the probability of response to vaccination (titre ≥0.17 IU/mL), and 2) variables that were associated with strength of protection given a response.

An intercept-only repeated measures multilevel (including village, farm and cow levels) linear mixed model was fitted using log VNA titre as an outcome variable to determine whether the village and farm level random effects needed to be included. Once it was confirmed that they did not, a generalised estimating equation (GEE) model (31) with a binomial error distribution was then used to model the probability of response (titre ≥0.17 IU/mL). Initially several univariable models were created, with age, BCS, sex, breed and herd size. For this analysis BCS was recoded as 2, 2.5 and ≥3, herd size as 1-6, 7-8 and 9-11, and age as ≤1.5-year old, 2-5 years old and ≥ 6-year old to create categories with even group sizes. Independent variables that were related (p ≤ 0.20) were then included in a multivariable GEE model. Backward elimination was then used until all the independent variables in the model had p ≤0.05, except that a confounder that changed coefficients or standard errors of independent variables by ≥20% was forced into the model, regardless of its p-value. Independent variables that had been removed were then added back into the model one by one, and retained if p ≤0.05. After building the main effect model, two-way interactions were created using all pairs of independent variables, interactions were included in the final model if p ≤0.05. This model building strategy was then used to build a GEE model of the strength of protection (log transformed titre). This model had first order auto-regressive work correlation matrix and normal error distribution. These analyses were carried out using STATA 13.1 (StataCorp, USA).

## Results

### Proportion of animals that responded to the vaccination

On day 0, all tested cattle had titres <0.17 IU/mL. Based on this cut-off point, the overall response rate for the intramuscular group was 98% and for the intradermal group 96% (Table 1). The overall proportion of the cattle with VNA titre ≥ 0.17 IU/mL between the two vaccination groups was not affected by route of vaccination (p=0.35) or the interaction between vaccination route and time after vaccination (p=0.31). However, compared to Day 0, the proportion of vaccinated cattle with a titre >0.17 IU/mL was greater on all other sample days (p<0.001); this was also the case for days 14 and 30 compared to day 90 (p≤0.02).

**Table 1.**
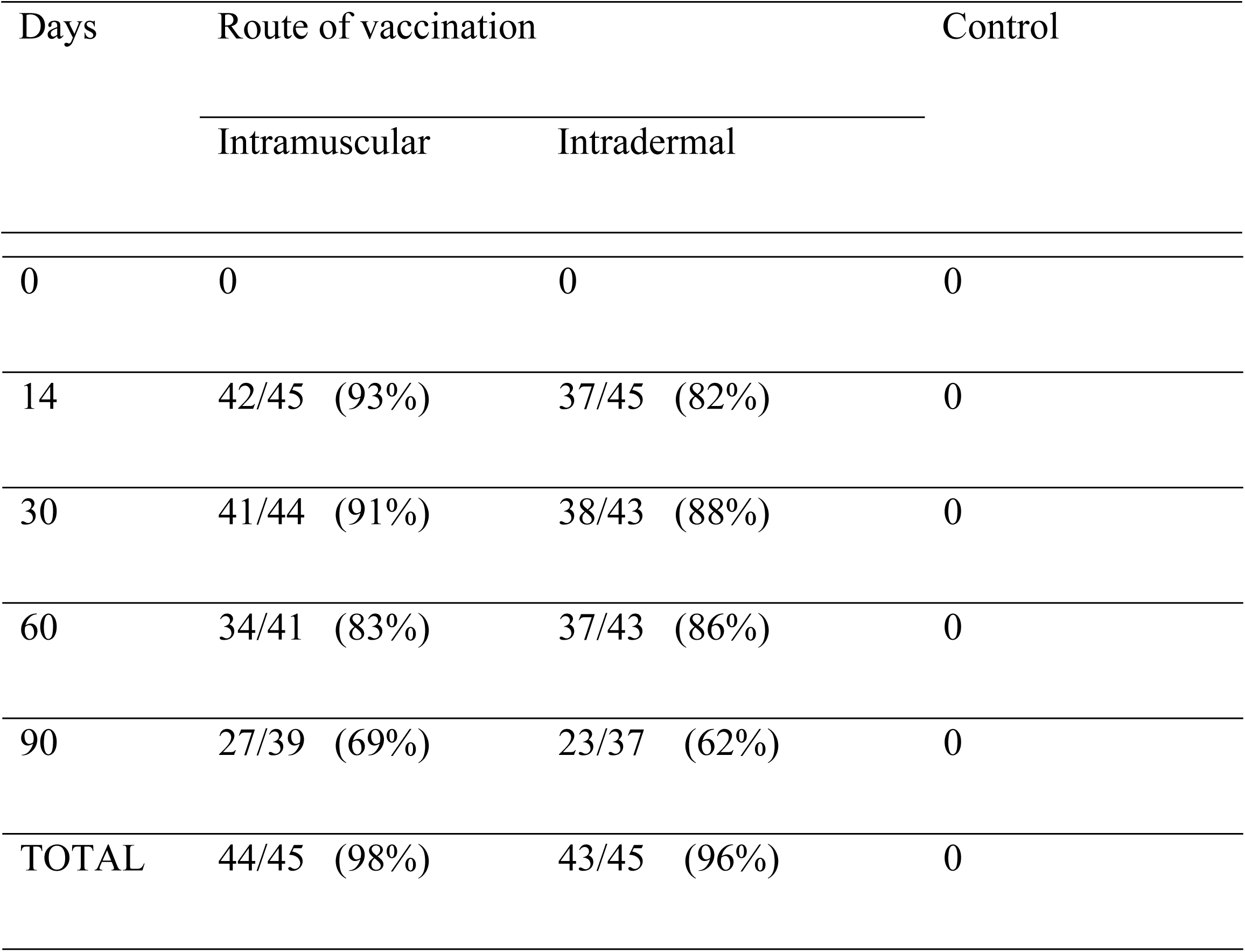
Proportion of cattle that responded to rabies vaccination (based on VNA titre ≥0.17 IU/mL)

| Days | Route of vaccination |  | Control |
| --- | --- | --- | --- |
|  | Intramuscular | Intradermal |  |
| 0 | 0 | 0 | 0 |
| 14 | 42/45 (93%) | 37/45 (82%) | 0 |
| 30 | 41/44 (91%) | 38/43 (88%) | 0 |
| 60 | 34/41 (83%) | 37/43 (86%) | 0 |
| 90 | 27/39 (69%) | 23/37 (62%) | 0 |
| TOTAL | 44/45 (98%) | 43/45 (96%) | 0 |

### Proportion of animals with protective VNA titres

The proportion of vaccinated cattle with the lower protective threshold (VNA titre ≥0.24 IU/mL) is summarised in Table 2.

**Table 2.** Proportion of cattle in each vaccination group with VNA titre ≥ 0.24 IU/mL.

| Days | Route of vaccination |  |
| --- | --- | --- |
|  | Intramuscular | Intradermal |
| 0 | 0 | 0 |
| 14 | 38/45 (84%) | 29/45 (64%) |
| 30 | 37/44 (84%) | 30/43 (70%) |
| 60 | 26/41 (63%) | 27/43 (63%) |
| 90 | 18/39 (46%) | 14/37 (39%) |
| TOTAL | 41/45 (91%) | 40/45 (89%) |

As with the proportion of cattle with a titre ≥0.17 IU/mL, there was no effect of vaccination route or interaction between vaccination route and time on the odds of a vaccinated animal having a titre ≥0.24 IU/mL (p= 0.67 and 0.38, respectively), but there was an effect of time since vaccination (p<0.001). Fig 1 depicts the effect of time since vaccination irrespective of vaccination routes.

**Fig 1.**
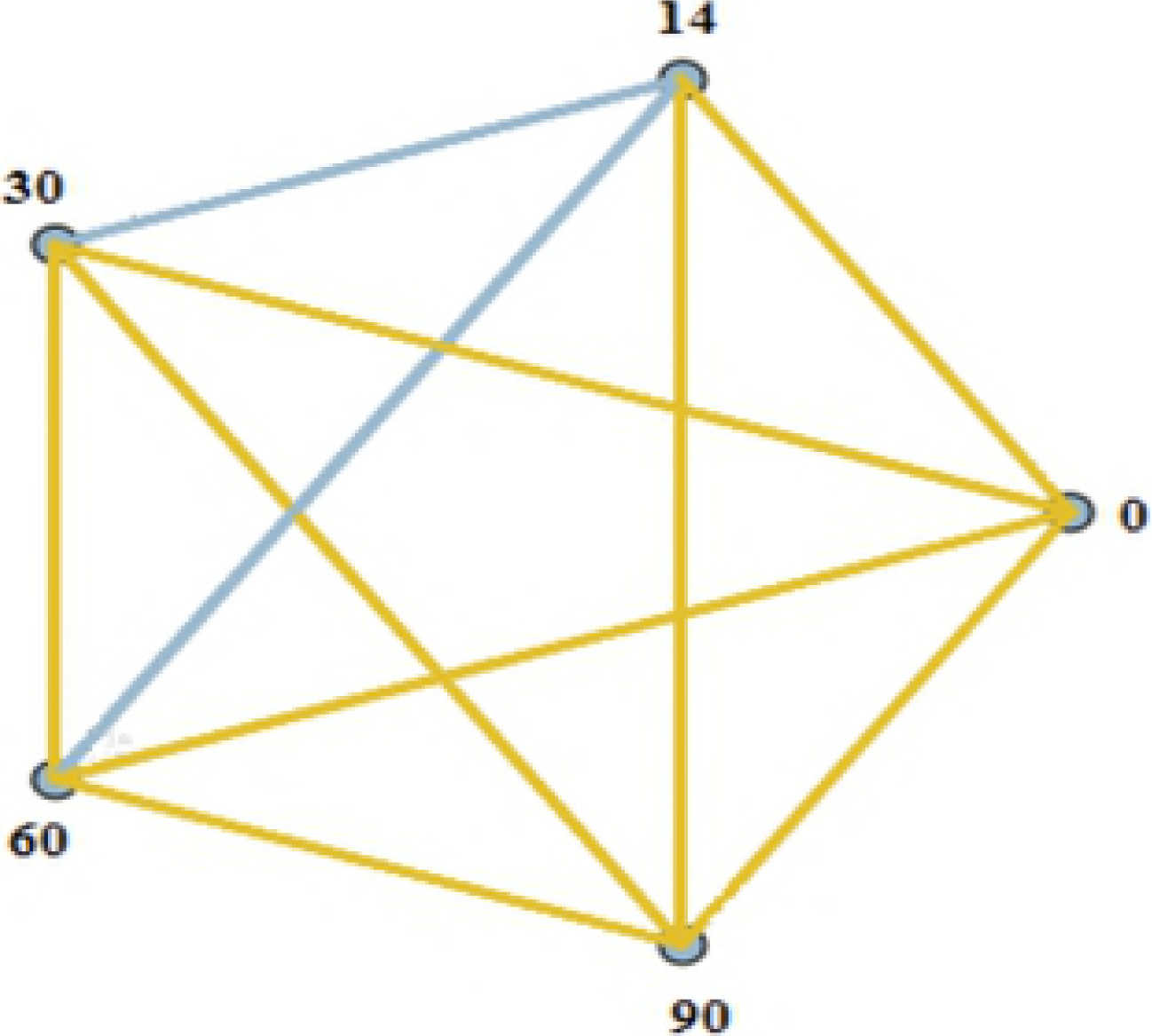
Pairwise comparisons of the effect of time since vaccination on proportion of vaccinated cattle with rabies VNA titres ≥0.24 IU/mL. Blue lines: Comparison between day 14 and day 30 (p=0.7), and comparison between day 14 and day 60 (p=0.07). Gold lines, p <0.002 for all comparisons, except day 60 and day 30 where p=0.02.

For the 0.5 IU/mL threshold, no effect of vaccination route on proportion of titres ≥0.5 IU/mL was found (p= 0.538). However, both the interaction between vaccination route and time and time alone were significant at the 4% level (p=0.039 and <0.001, respectively). On both day 14 and day 30 the proportion of cattle with a VNA titre ≥ 0.5 IU/mL was lower in the intradermally vaccinated group than in the intramuscularly vaccinated group (Table 3).

**Table 3:** Proportion of cattle with VNA titre ≥ 0.5 IU/mL at 0,14,30,60 and 90 days.

| Days | Route of vaccination |  | Comparison between vaccination groups |
| --- | --- | --- | --- |
|  | Intramuscular | Intradermal |  |
| 0 | 0 | 0 | - |
| 14 | 35/45 (78%) | 16/45 (36%) | $p < 0.001$ |
| 30 | 34/44 (77%) | 25/43 (58%) | $p = 0.02$ |
| 60 | 20/41 (49%) | 18/43 (42%) | $p = 0.4$ |
| 90 | 11/39 (28%) | 9/37 (24%) | p=0.7 |
| TOTAL | 40/45 (89%) | 32/45 (71%) |  |

### Effect of vaccination route and time on VNA titres

Time, vaccination route and their interaction were found to have an effect on VNA titres (p<0.001) (see Fig 2). In both groups, VNA titres were higher after vaccination throughout the study period than before vaccination (p<0.001 for all comparisons), with the mean VNA titres of both groups peaking on day 30. The decline in titres after the peak was more marked in the intramuscularly vaccinated group than the intradermal group with titres being lower on day 60 than on day 30 in the former group (p<0.001), but only by day 90 in the intradermally vaccinated group.

**Fig. 2.**
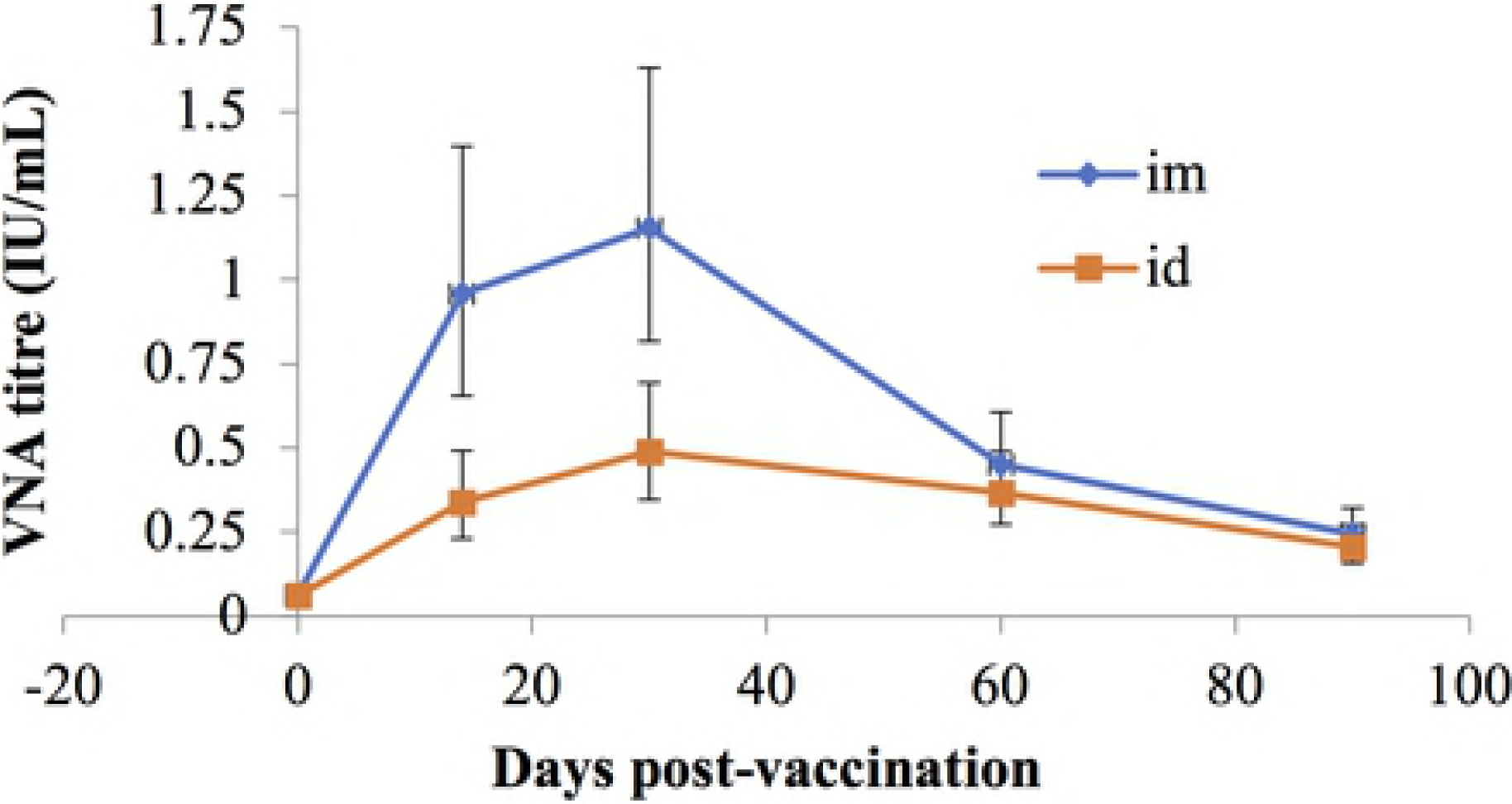
Geometric mean VNA titres of intramuscularly (im) and intradermally (id) vaccinated cattle on 0, 14, 30, 60 and 90 days post vaccination.

Mean antibody titres were lower on days 14 and 30 in the intradermally vaccinated group than in the intramuscularly vaccinated group. On day 14 the back transformed mean difference between the VNA titres of intramuscularly and intradermally vaccinated animals was 0.62 (95% CI 0.02 to 1.3) IU/mL, whereas on day 30 it was 0.66 (95% CI 0.22 to 1.39) IU/mL. Thus, in terms of the antibody response, the intradermal vaccination was inferior to the intramuscular vaccination.

### Effect of cow and farm level factors on response to vaccination

The GEE modelling process found that there was a significant interaction between route and age on the likelihood of a sample having a titre ≥0.17 IU/mL. For cattle <2 years old the odds of a sample titre being ≥0.17 IU/mL; was 4.8 (95%CI: 1.3-17.7) times greater for intradermal than intramuscular vaccinated animals group, whereas for animals between 2 and 5 years of age the equivalent odds ratio was 0.24 (95%CI: 0.06-0.97), and for cattle >5 years old 0.34 (95%CI: 0.04-2.73). Apart from time the only factor which influenced the odds of an antibody response ≥0.17 IU/mL, was herd size, with samples from herds with more than six cattle having lower odds of titres ≥0.17 IU/mL.

For the actual titres, apart from time, the only significant effect found was an effect of age and its interaction with route of vaccination. For intramuscularly vaccinated cattle, mean titre increased with age whereas for intradermally vaccinated cattle it decreased. Titres of 2-year-old cattle vaccinated intradermally were higher than those of 2-year-old cattle vaccinated intramuscularly; this was reversed in the other two age categories.

## Discussion

This is the first study to compare the efficacy of intramuscular and intradermal routes of rabies vaccination in cattle in Bhutan under field conditions. It is also, as far as the authors are aware, the first study of intradermal vaccination against rabies in cattle to use more than 10 cattle per treatment group. The study was designed as a non-inferiority trial with the aim of confirming whether the mean VNA titres produced by intradermal vaccination were no more than 0.5 IU/mL lower than the titres produced by the standard intramuscular vaccination. In addition, three thresholds of vaccination response were used in order to further compare the response of the two vaccination routes.

The geometric mean VNA production by intradermal vaccination using 1/5 (0.2mL) of the dose used in standard intramuscular (1mL) route was significantly lower than the standard intramuscular route on days 14 and 30 post vaccination. The back transformed mean difference between intramuscular and intradermal groups was >0.6 IU/mL, indicating that based on the criteria of the study, intradermal vaccination was inferior to intramuscular vaccination. Furthermore, the geometric mean titre in the intradermally vaccinated cattle did not achieve the WHO and OIE recommended threshold titre of ≥0.5 IU/mL on any day post vaccination. However, overall 71% (32/45) of the intradermally vaccinated cattle had a titre ≥0.5 IU/mL on at least one day. These proportions were significantly lower (P<0.02) than the intramuscular group only on days 14 and 30 post vaccination - with 36 and 58% having titres ≥0.5 IU/mL on day 14 and 30, respectively compared to the equivalent figures of 78 and 77% in the intramuscular group.

This finding is in contrast to the findings of Asokkumar et al. (19) that used the same vaccine brand as used in this study. The VNA titres were measured using Rapid Fluorescent Focus Inhibition Test (RFFIT). They reported intradermal vaccination produced titres equivalent to those produced by the intramuscular route despite using 1/5 of the dose (VNA titre in IU/mL; Zero day- 0.07 for IM and ID, 14^th^ day 0.89 for IM and 0.6 for ID; 28^th^ day 2.81 for IM and 2.54 for ID). However, in addition to being a small study (8 cattle per treatment group), this was a post-exposure prophylaxis study, so cattle were vaccinated on days 0,3,7,14 and 28, significantly increasing the chance of a response. Furthermore, as a post-exposure study with no untreated controls, it is not clear whether any of the response was due to exposure to wild-type virus.

A more directly comparable study is that by Benisek et al. (2006) (20) who in unexposed cattle reported that the VNA response in their intradermally vaccinated group was significantly greater than intramuscular group despite using only 1/5^th^ of the dose in the former group. The VNA titres, measured by RFFIT, for day 14 were 0.51 IU/mL and 0.73 IU/mL for intramuscular and intradermal routes respectively. Similarly, the day 35 and 90 were also greater for the intradermal group than the intramuscular (1.64 IU/mL vs 1.07 IU/mL and 1.40 IU/mL vs 0.94 IU/mL respectively).

It is unclear why Benisek et al. (20) found different results from this study. Although they used a different brand of vaccine (Rabicell) it was also an inactivated rabies vaccine with an aluminium hydroxide adjuvant. The response to the intramuscular vaccination reported by Benisek et al. (20) was different from that seen in this study. The mean VNA titres in their study were still >0.5 IU/mL 180 days after intramuscular vaccination compared to this study in which mean VNA titres after intramuscular vaccination were <0.5 IU/mL within 90 days. Furthermore, the proportion of cattle with a titre ≥0.5 IU/mL on days 14 and 30 in the intramuscularly vaccinated group in this study were lower than the WHO targets for tissue culture rabies vaccine of almost 100% (32).

The results of this study also seem to be in contrast to the undoubted efficacy of intradermal rabies vaccination in humans (17). However, in humans pre-exposure vaccination is a multidose regimen that requires three to four doses of vaccine (33, 34). Furthermore, the results of the current study are consistent with the statement made by WHO (30) that ‘antibody titres are higher and more sustained after intramuscular injection’.

One potential issue is that intradermal vaccination is more difficult, so some vaccines could have been incorrectly administered into adipose or subcutaneous tissue. However, >90% of vaccinations were recorded as definitively going intradermally, and the titres of the nine cattle in which this was not the case were not significantly different from those which did (data not shown). Another potential issue was that as there was no licensed rabies vaccine for intradermal use in cattle, a 10 mL vaccine vial was used for this study. Repeated drawing of vaccine from this multidose vial could have resulted in some animals receiving a dose less than 0.2 mL. Finally, cattle were released for grazing after vaccination and were not monitored afterwards. As intradermal administration can cause irritation at the vaccination site (35), rubbing induced by irritation at the injection site could have led to leakage of vaccine before being absorbed into the system. Thus, it is plausible that despite a sufficient dose being given intradermally, the vaccine was not retained long enough in the dermal tissue to be absorbed.

Nonetheless, all of these issues would affect individual cows, thereby increasing variability in VNA response between animals not vaccinated correctly and those which were. However, there was no evidence of variability in this study, in contrast to Benisek et al. (20) study in which intradermal vaccination was associated with an increase in variability of VNA titres. Thus, the most plausible rationale for the difference between the results of this study and that of Benisek et al. (20) is differences between vaccines, different tests for VNA measurements (RFFIT vs FAVN test) or between the cattle in each study. The data from this suggest that one of the most important of these is that Benisek et al. (20) used young bulls in their study while cattle of all ages were used in this study. Furthermore, a classical antibody response curve was produced in this study with a rise on day 14^th^ after vaccination, peak at day 30 and fall on day 60 and 90. However, this titre curve was not observed in the Benisek et al. (20) study.

In this study there was a significant interaction between vaccination and age with younger animals (age <2) having an increased chance of response (titre ≥0.17 IU/mL) when given intradermal rather than intramuscular vaccination and having higher titres when they responded. Further research is required to better establish whether the apparent association between age and response to intradermal vaccination is clinically significant.

Even disregarding the impact of age the effect of route on VNA titres may not have been as great as suggested in Figure 2. When the cut-off for a protective titre was reduced from ≥0.5 to ≥0.24 IU/mL, there was no difference in response between the two routes and geometric mean VNA titres in both groups were ≥0.24 IU/mL throughout the duration of the study following vaccination. This cut-off point was chosen based on the vaccination and challenge studies conducted in cattle, dogs, cats (27, 28, 36, 37) and foxes (29). Côrtes et al. (28) reported a protection rate of 80% in cattle with titre ≥0.24 IU/mL when challenged with a virulent strain of rabies virus. However, there was only a 5% increase in the proportion protected with titre ≥ 0.5 IU/mL.

Thus, although a higher VNA titre is preferred, any seroconversion following vaccination indicates some degree of protection (27, 37), particularly against natural infection, which is usually less severe than the experimental infection used to set thresholds (27, 38). However, much of this is based on data from dogs, which are reservoir hosts and there may therefore be a certain degree of host-virus adaptation that reduces the risk of infection in dogs compared to dead end hosts such as cattle (27).

Thus, the results of this study do not support the routine use of intradermal vaccination of cattle using the Raksharab vaccine at a dose rate of 0.2 mL, except, perhaps in cattle <2 years old. Further studies are needed, in particular on the use of a booster vaccination 60 days after primary vaccination, or on the use of a higher dose than the 0.2 IU/mL used in this study. The latter is likely to have lower costs and be more feasible, and increasing the intradermal dose is easier in cattle than humans as cattle skin is relatively thick and it is therefore easier to administer larger quantities of vaccine at one site (39).

## Conclusion

Intradermal vaccination induced protective titres (≥ 0.5 IU/mL) in 71% of cattle, despite using 1/5 of the recommended dose. However, the proportion of cattle with a protective titre was significantly lower than for cattle given the standard dose (1mL) intramuscularly (71 vs 89%). In addition, mean antibody titres in the intradermally vaccinated cattle were significantly lower than the intramuscularly vaccinated cattle on days 14 and 30 post vaccination. Intradermal vaccination using 1/5 dose was inferior to intramuscular vaccination using the standard dose. However, the antibody responses obtained in this study were good enough to support further testing of intradermal vaccination with an increased dose.

## Acknowledgement

We would like to sincerely acknowledge Alexandre Servat, Valère Brogat, Sébastien Kempf, Anouck Labadie, Estelle Litaize, Jonathan Rieder, Laetitia Tribout and Jean Luc Schereffer at ANSES, Nancy Laboratory for Rabies and Wildlife, France for analysing the rabies vaccine and serum samples.

We are grateful to Dr. Sonam Peldon and her team at Haa, Bhutan for helping to collect and process samples.

We are also thankful to all the cattle owners who agreed to participate in this study, the New Zealand Development Aid for providing this scholarship, the Royal Government of Bhutan and Department of Livestock for giving the authorizations to conduct this study in Bhutan.

Lastly, we thank Kencho Sum (Triple Gem), my family members and friends (Xue Qi Soon, Dr. Mary Gaddam, Dr. Linda Laven) for giving us the moral support and inspiration to pursue and complete this study.

## Author contribution

Conception, study design and project administration: KW. Sample analysis: FC, MW. Provision of resources: KW, RL, FC, MW. Data analysis and interpretation: RL, AY, KW. Paper writing: KW, RL, FC, MW, AY.

